## Supplementary figure 1-12 and Supplementary table 1 for "MiDAS 5: Global diversity of bacteria and archaea in anaerobic digesters"

[1]Center for Microbial Communities, Department of Chemistry and Bioscience, Aalborg University, Aalborg, Denmark. [2]Centre of Biological Engineering, University of Minho, Portugal. [3]School of Water, Energy and Environment, Cranfield University. Cranfield, United Kingdom. [4]Australian Centre for Water and Environmental Biotechnology (ACWEB), The University of Queensland, Australia. [5]Department of Civil and Environmental Engineering, University of Massachusetts Amherst, MA, USA. [6]National University of Salta, Salta, Argentina. [7]Department of Chemical Engineering, Lund University, Sweden. [8]INGEBI-CONICET, University of Buenos Aires, Argentina. [9]Laboratory for Environmental Biotechnology, Ecole Polytechnique Fédérale de Lausanne (EPFL), Switzerland. [10]Chair of Urban Water Systems Engineering, Technical University of Munich (TUM), Garching, Germany. [11]Institute of Water Quality and Resource Management, TU Wien, Austria. [12]Department of Urban and Environmental Engineering & Graduate School of Carbon Neutrality, Ulsan National Institute of Science and Technology (UNIST), South Korea. [13]School of Chemical Engineering, National Technical University of Athens, Greece. [14]Applied Environmental Biotechnology Laboratory, Birla Institute of Technology and Science (BITS-Pilani), India. [15]School of Biological and Chemical Sciences and Ryan Institute, University of Galway, Ireland. [16]Water Supply and Bioeconomy Division, Faculty of Environmental Engineering and Energy, Poznan University of Technology, Poland. [17]Department of Water Technology and Environmental Engineering, University of Chemistry and Technology Prague, Czech Republic. [18]Research Scientist at Kemira Oyj, Espoo R&D Center, Finland. [19]Chemical Engineering Department, Khalifa University, United Arab Emirates. [20]Environmental Science and Engineering Program, Biological and Environmental Science and Engineering Division, King Abdullah University of Science and Technology (KAUST), Saudi Arabia. [21]Center for Microbial Ecology and Technology (CMET), Ghent University, Belgium.

*Correspondence to: Per Halkjær Nielsen, Center for Microbial Communities, Department of Chemistry and Bioscience, Aalborg University, Fredrik Bajers Vej 7H, 9220 Aalborg, Denmark; Phone: +45 2173 5089; Fax: Not available; or Morten Kam Dahl Dueholm, Center for Microbial Communities, Department of Chemistry and Bioscience, Aalborg University, Fredrik Bajers Vej 7H, 9220 Aalborg, Denmark; Phone: (+45) 9940 8508; Fax: Not available;

**Running title:** Global microbiota of anaerobic digesters

**Table of content:**

Page 3-14: Supplementary Fig. 1-12
Page 15: Supplementary Table 1
Page 16: Supplementary References

### Supplementary Figures:

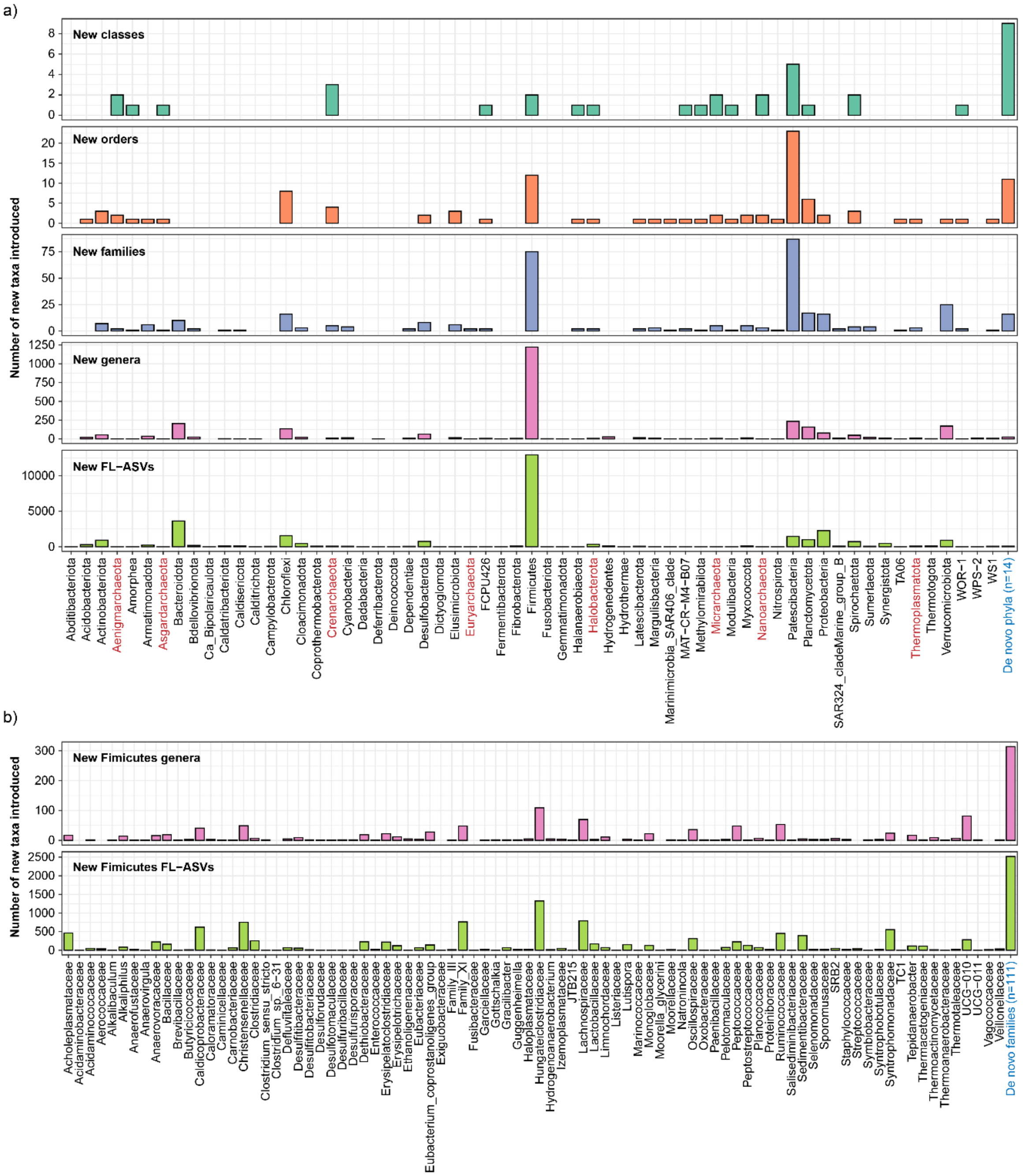


Supplementary Fig. 1: Taxonomic affiliation of new reference sequences (FL-ASVs) added to the MiDAS database in this study. a) Phylum level distribution of new FL-ASVs introduced. Archaeal phyla are highlighted with red labels. b) Family level distribution of new FL-ASVs belonging to the Firmicutes. All de novo phyla or families were merged into one category highlighted with blue labels. The numbers indicate how many taxa were merged.

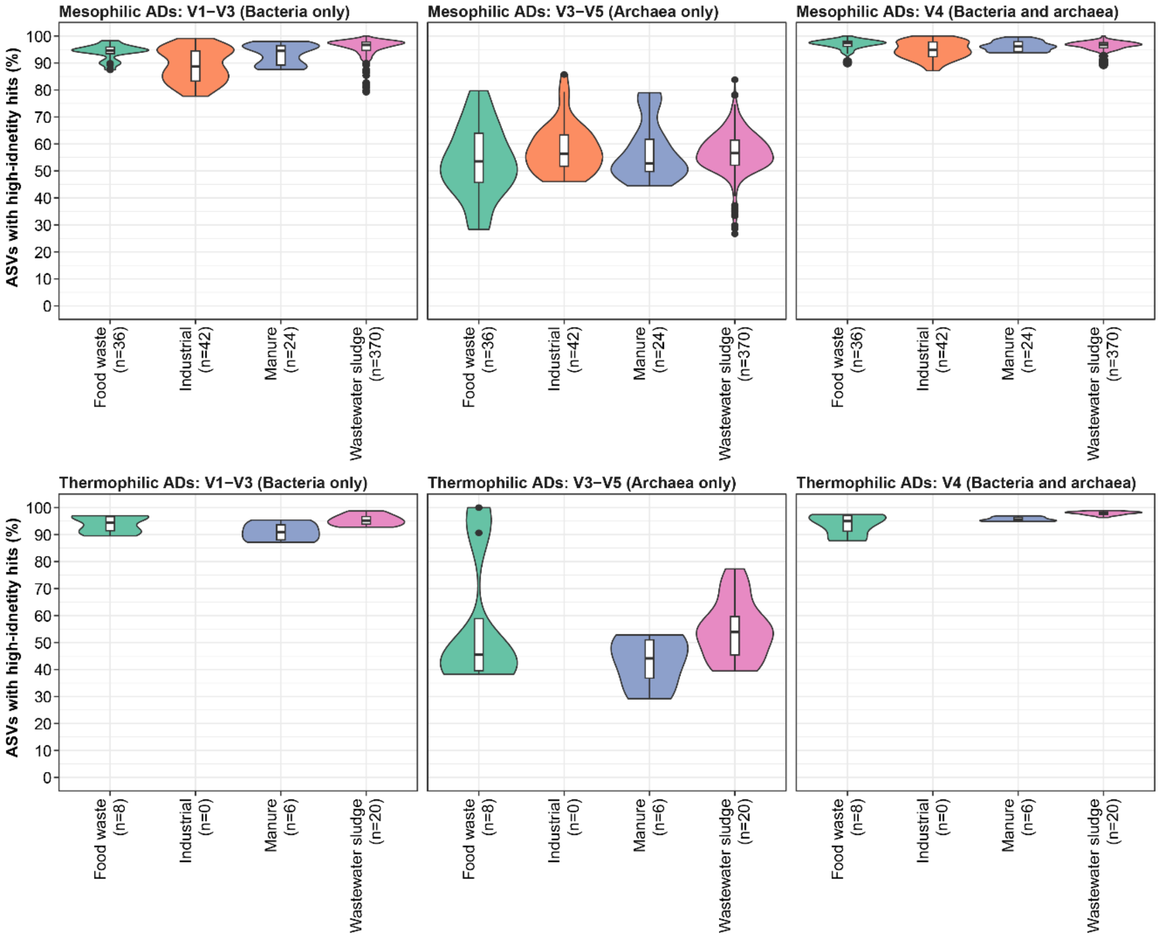


Supplementary Fig. 2: Database coverage for mesophilic and thermophilic ADs treating different primary substrates based on short-amplicon data from this study. The ASVs for each of the samples were filtered based on their relative abundance (only ASVs with ≥0.01% relative abundance were kept) before the analyses. The percentage of the microbial community represented by the remaining ASVs after the filtering was 95.29% ± 2.24% (mean±SD) for V1-V3 amplicons (only bacteria), 99.64% ± 0.17% for V3-V5 amplicons (mainly archaea), and 97.32% ± 2.07% for V4 amplicons (bacteria and archaea) across the number of duplicate biological independent samples (n) indicated on the x-axis. High-identity (≥99%) hits were determined by stringent mapping of ASVs to the MiDAS 5.2 database. The violin and box plot represent the distribution of percent of ASVs with high-identity hits for each combination of primary substrate and temperature (mesophilic or thermophilic). Box plots indicate median (middle line), 25th, 75th percentile (box), and the min and max values after removing outliers based on 1.5x interquartile range (whiskers). Different colors are used to distinguish the different primary substrates.

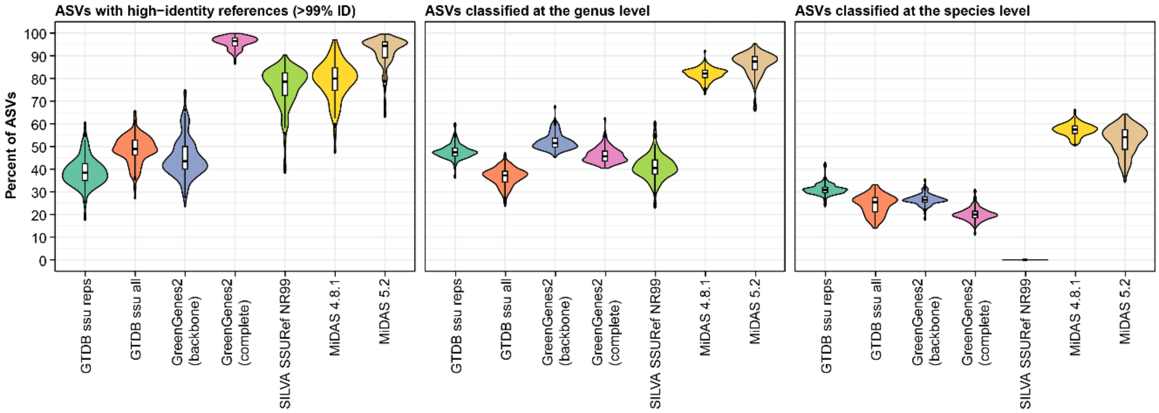


Supplementary Fig. 3: Database evaluation based on short-amplicon data from Mei *et al.* 2017 [1]. The ASVs for each of the samples were filtered based on their relative abundance (only ASVs with ≥0.01% relative abundance were kept) before the analyses. The percentage of the microbial community represented by the remaining ASVs after the filtering was 95.00% ± 1.84%. High-identity (≥99%) hits were determined by stringent mapping of ASVs to each reference database. Classification of ASVs was done using the SINTAX classifier. The violin and box plot represent the distribution of percent of ASVs with high-identity hits or genus/species-level classifications for each database across n=194 biologically independent samples. Box plots indicate median (middle line), 25th, 75th percentile (box), and the min and max values after removing outliers based on 1.5x interquartile range (whiskers). Outliers have been removed from the box plots to ease visualization. Different colors are used to distinguish the different databases: GTDB_bac120_ssu_reps_r214, GTDB_ssu_all_r214, GreenGenes2_2022_10 (backbone and complete database), SILVA 138.1 SSURef NR99, MiDAS 4.8.1, and MiDAS 5.2.

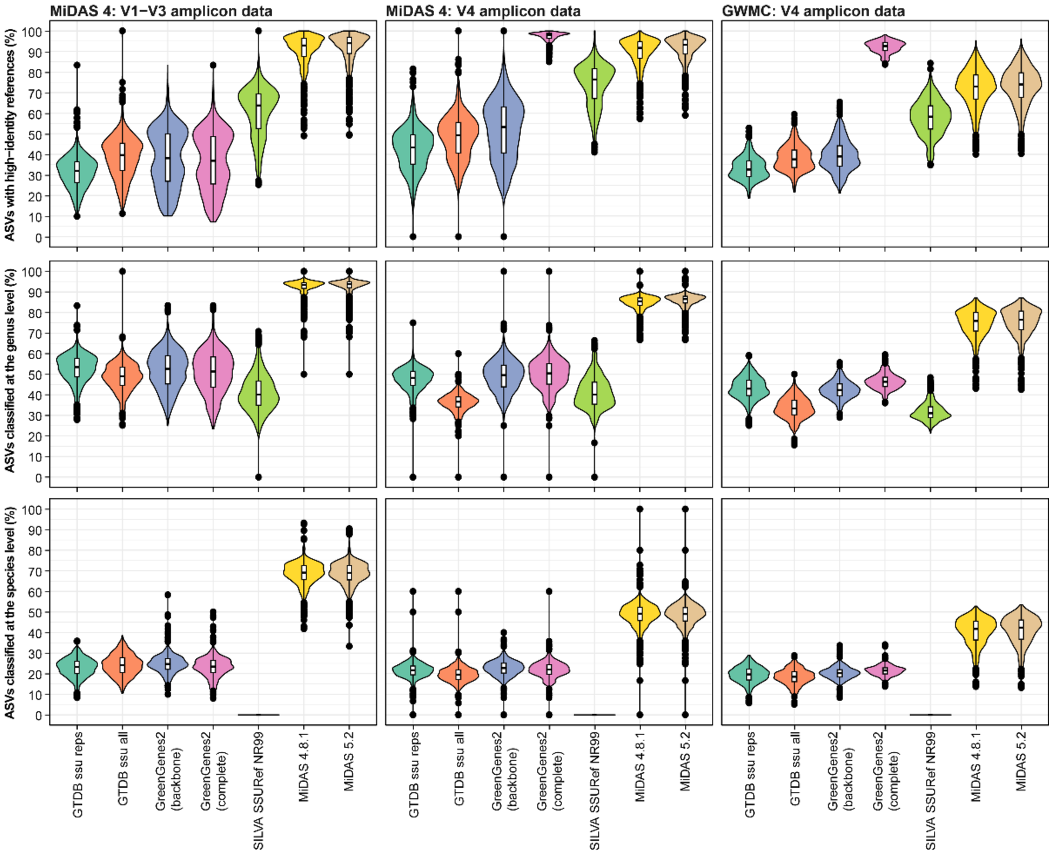


Supplementary Fig. 4: Database evaluation based on short-amplicon data from the MiDAS Global WWTP project (MiDAS 4) [2] and the Global Wastewater Microbiome Consortium project (GWMC) [3]. The ASVs for each of the samples were filtered based on their relative abundance (only ASVs with ≥0.01% relative abundance were kept) before the analyses. The percentage of the microbial community represented by the remaining ASVs after the filtering was 92.69% ± 3.22% (mean±SD) for MiDAS 4 V1-V3 amplicons, 96.4% ± 2.27% for MiDAS 4 V4 amplicons, and 88.35% ± 2.98% for GWMC V4 amplicons across samples. High-identity (≥99%) hits were determined by stringent mapping of ASVs to each reference database. Classification of ASVs was done using the SINTAX classifier. The violin and box plot represent the distribution of percent of ASVs with high-identity hits or genus/species-level classifications for each database across n=1279 (MiDAS 4 V1-V3), n=1278 (MiDAS 4 V4), and n=1165 (GWMC V4), biologically independent samples duplicates. Box plots indicate median (middle line), 25th, 75th percentile (box), and the min and max values after removing outliers based on 1.5x interquartile range (whiskers). Outliers have been removed from the box plots to ease visualization. Different colors are used to distinguish the different databases: GTDB_bac120_ssu_reps_r214, GTDB_ssu_all_r214, GreenGenes2_2022_10 (backbone and complete database), SILVA 138.1 SSURef NR99, MiDAS 4.8.1, and MiDAS 5.2.

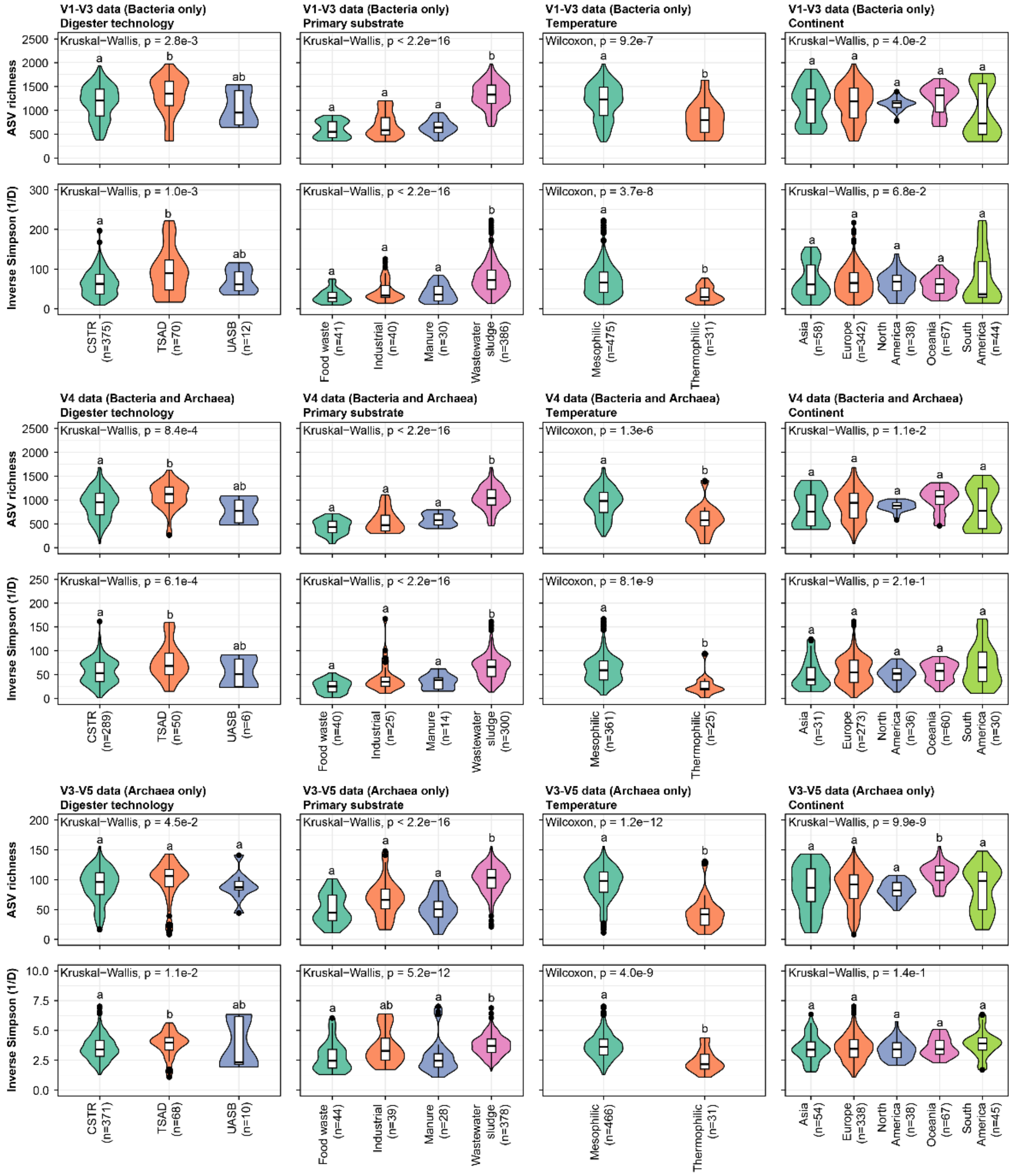


Supplementary Fig. 5: Effect of process parameters and geography on alpha diversity. The non-parametric Kruskal-Wallis test was used to determine the statistical support for differences between the means of the groupings in the V1-V3, V3-V5, and V4 amplicon data set. The exact value could not be determined for $p < 2.2*10^{-16}$. A post-hoc Dunn's test (Bonferroni correction, α=0.01) was used for pairwise comparison of individual groups and the results are shown with compact letter display (groups that do not share letters are significantly different). The violin and box plot represent the distribution of ASV richness or inverse Simpson (1/D) across the number of biological independent samples (n) indicated on the x-axis. The box plots indicate median (middle line), 25th, 75th percentile (box), and the min and max values after removing outliers based on 1.5x interquartile range (whiskers). Outliers are shown as black dots. CSTR: Continuous stirred-tank reactor; TSAD: Two-stage anaerobic digestion; UASB: Upflow anaerobic sludge blanket. Different colors are used to distinguish the categories for each examined variable.

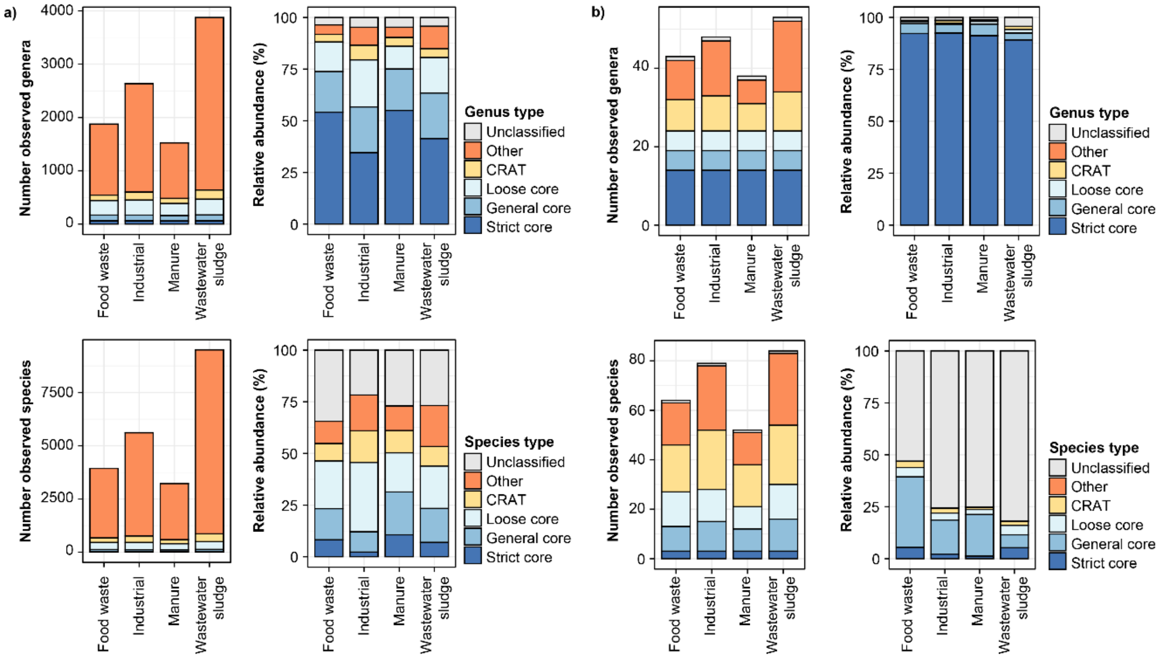


Supplementary Fig. 6: Core and conditionally rare or abundant taxa in anaerobic digesters globally. a) and b) Number of observed genera (top) and species (bottom), respectively, and their relative abundance in mesophilic ADs treating different primary substrates based on a) V1-V3 (Bacteria only) and b) V3-V5 (Archaea only) amplicon data. Values for genera and species are divided into strict core, general core, loose core, CRAT, other taxa, and unclassified ASVs based on the "most wanted" list (Supplementary Data 3). The relative abundance of different groups was calculated based on the mean relative abundance of individual genera or species across samples.

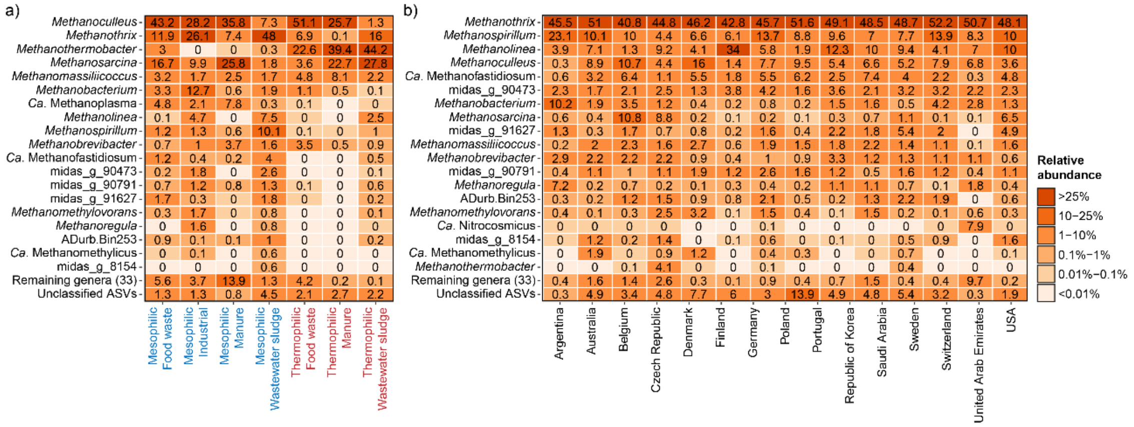


Supplementary Fig. 7: Top 25 archaeal genera based on V3-V5 amplicon data. The percent relative abundance represents the mean abundance relative to all archaea across a) different temperature range and primary substrates, and b) different countries considering only mesophilic ADs treating mainly wastewater sludge.

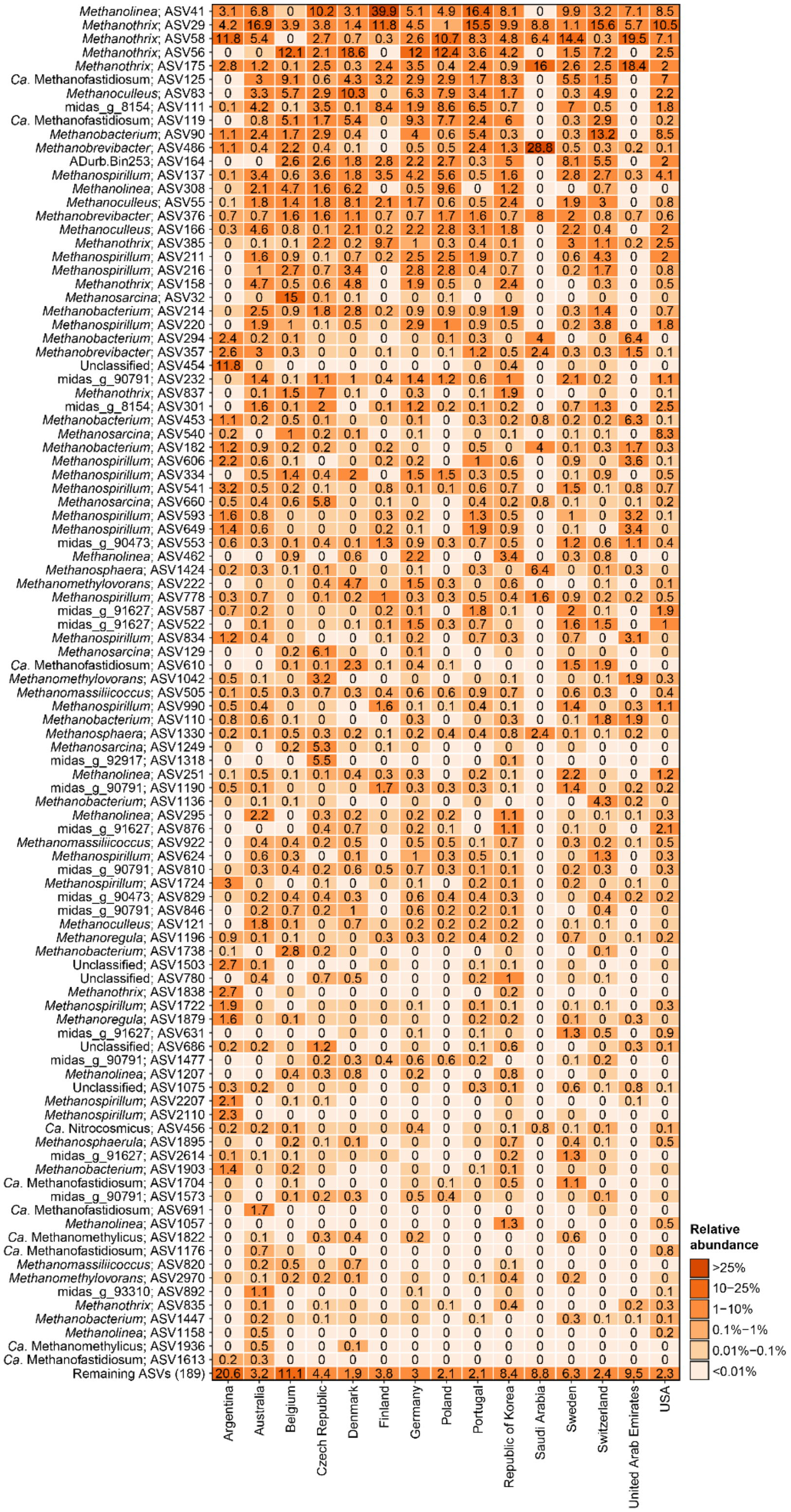


Supplementary Fig. 8: Top 100 archaeal ASVs based on V4 amplicon data. The percent relative abundance represents the mean abundance relative to all archaea across different countries considering only mesophilic ADs treating mainly wastewater sludge.

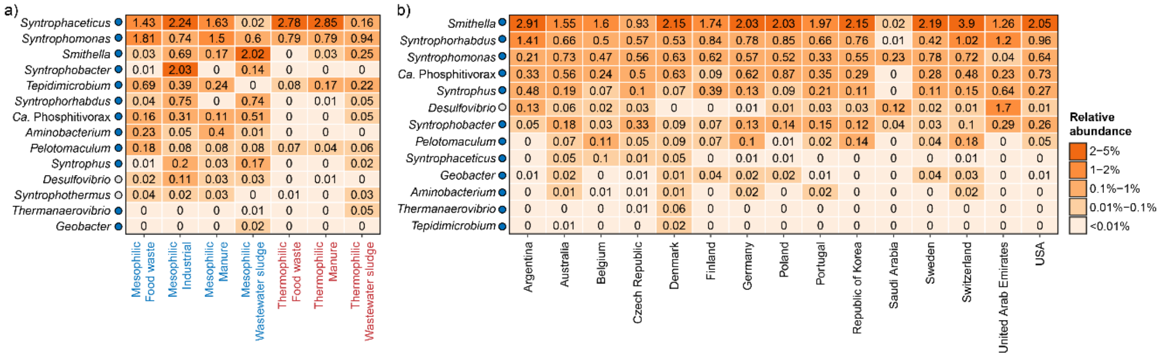

Supplementary Fig. 9: Global diversity of syntrophs based on V4 amplicon data. The percent relative abundance represents the mean for genera across a) different temperature range and primary substrates, and b) different countries considering only mesophilic ADs treating mainly wastewater sludge. Colored circles next to the genus labels indicate whether the genera have previously been identified as growing in ADs at Danish WWTPs according to Jiang *et al.* [4]. Blue: >50% of ASVs classified as growing; Yellow: 20-50% of ASVs classified as growing. Red: <20% of ASVs classified as growing. Gray: No information available for the specific genus.

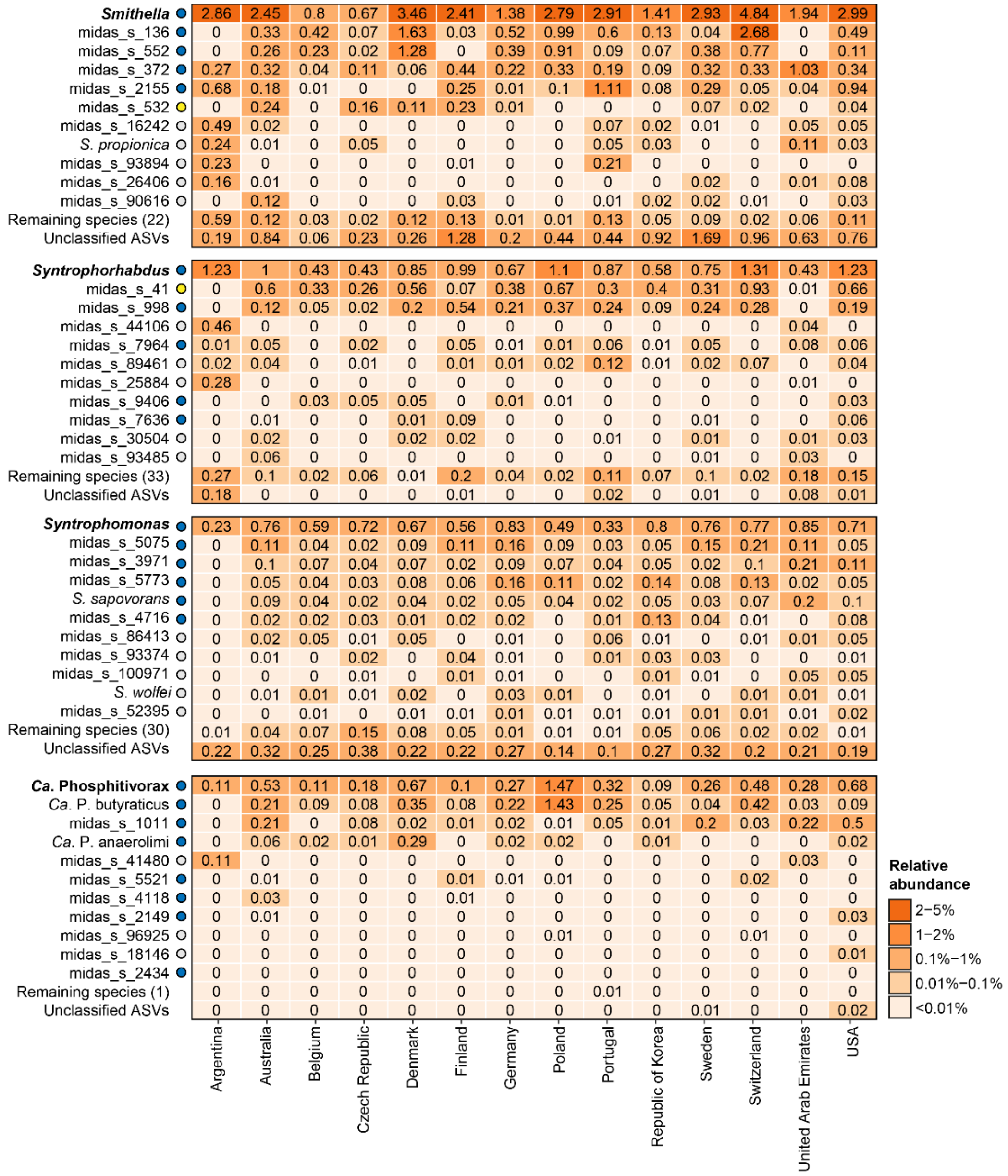


Supplementary Fig. 10: Species-level diversity of selected syntrophs based on V1-V3 amplicon data. The percent relative abundance represents the mean across different countries considering only mesophilic ADs treating mainly wastewater sludge. Colored circles next to the genus/species labels indicate whether the taxa have previously been identified as growing in ADs at Danish WWTPs according to Jiang *et al.* [4]. Blue: >50% of ASVs classified as growing; Yellow: 20-50% of ASVs classified as growing. Red: <20% of ASVs classified as growing. Gray: No information available for the specific taxa.

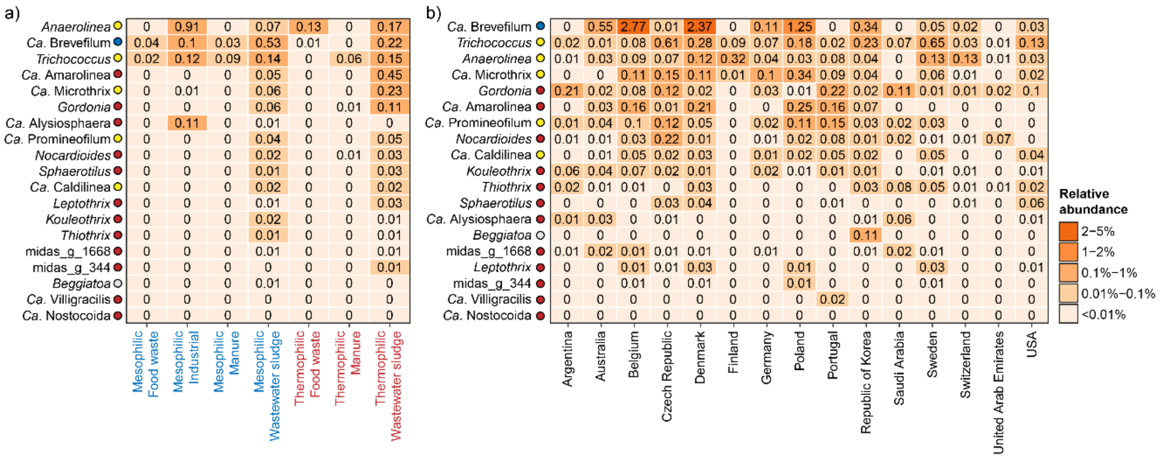


Supplementary Fig. 11: Global diversity of known filamentous organisms based on V4 amplicon data. The percent relative abundance represents the mean for genera across a) different temperature range and primary substrates, and b) different countries considering only mesophilic ADs treating mainly wastewater sludge. Colored circles next to the genus labels indicate whether the genera have previously been identified as growing in ADs at Danish WWTPs according to Jiang *et al.* [4]. Blue: >50% of ASVs classified as growing; Yellow: 20-50% of ASVs classified as growing. Red: <20% of ASVs classified as growing. Gray: No information available for the specific genus.

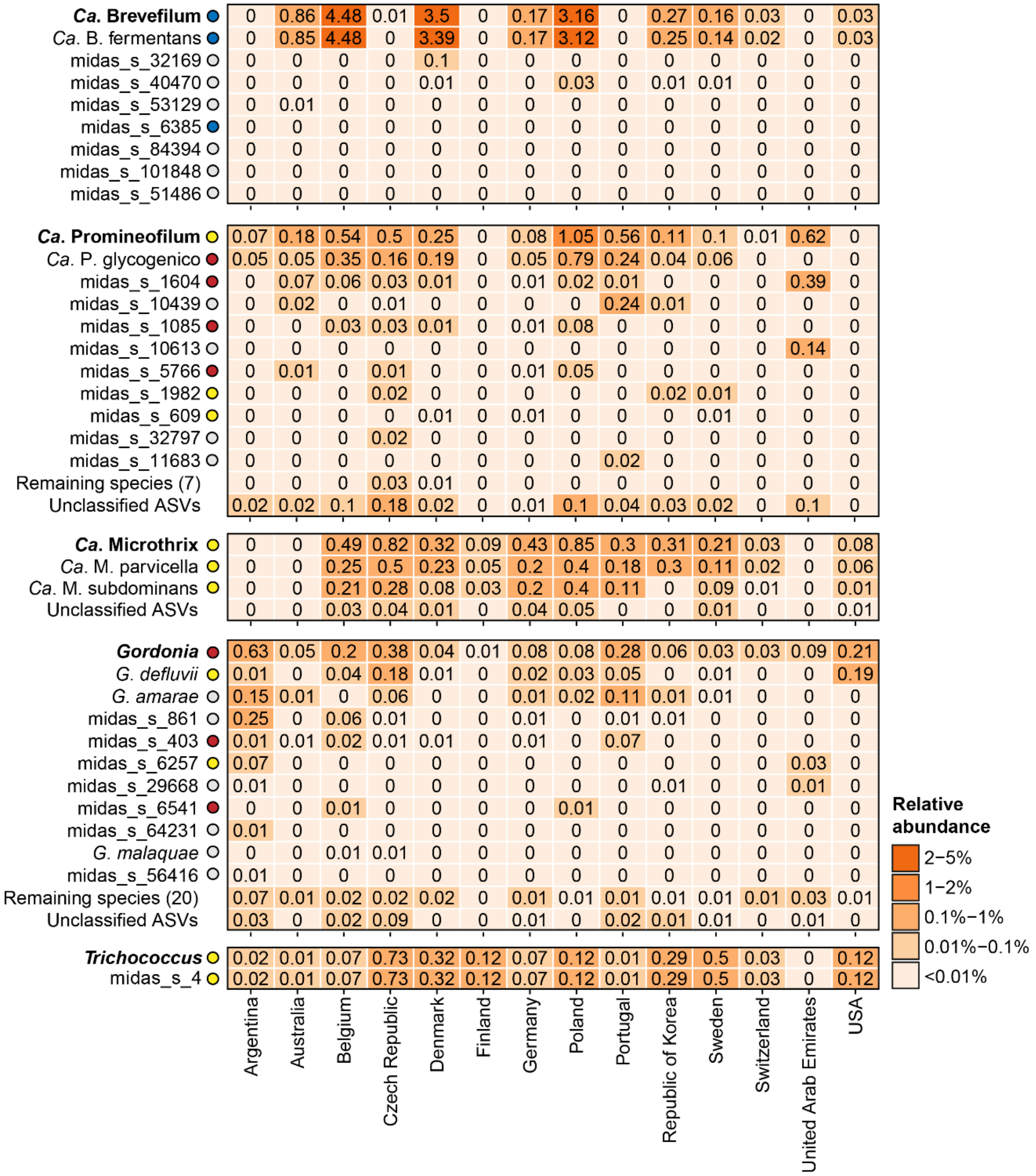


Supplementary Fig. 12: Species-level diversity of selected filaments based on V1-V3 amplicon data. The percent relative abundance represents the mean across different countries considering only mesophilic ADs treating mainly wastewater sludge. Colored circles next to the genus/species labels indicate whether the taxa have previously been identified as growing in ADs at Danish WWTPs according to Jiang *et al.* [4]. Blue: >50% of ASVs classified as growing; Yellow: 20-50% of ASVs classified as growing. Red: <20% of ASVs classified as growing. Gray: No information available for the specific taxa.

### Supplementary Tables:

Supplementary Table 1: Oligonucleotides used for full-length 16S rRNA gene library preparation. Unique molecular tags are marked in red. Primers specific for bacteria or archaea are marked with "_b_" or "_a_" in their names, respectively.

| **Name** | **Sequence (5' to 3')** |
|---|---|
| f16S_b_pcr1_fw | CAAGCAGAAGACGGCATACGAGATNNNYRNNNYRNNNYRNNNAGRGTTYGATYMTGGCTCAG |
| f16S_b_pcr1_rv | AATGATACGGCGACCACCGAGATCNNNYRNNNYRNNNYRNNNGACGGGCGGTGWGTRCA |
| f16S_a_pcr1_fw | CAAGCAGAAGACGGCATACGAGATNNNYRNNNYRNNNYRNNNTCCGGTTGATCCYGCBRG |
| f16S_a_pcr1_rv | AATGATACGGCGACCACCGAGATCNNNYRNNNYRNNNYRNNNGGCCATGCAMYWCCTCTC |
| f16S_pcr2_fw | CAAGCAGAAGACGGCATACGAGAT |
| f16S_pcr2_rv | AATGATACGGCGACCACCGAGATC |

**Supplementary references:**


1. Mei, R. *et al.* Operation-driven heterogeneity and overlooked feed-associated populations in global anaerobic digester microbiome. *Water Research* **124**, 77–84 (2017).
2. Dueholm, M. K. D. *et al.* MiDAS 4: A global catalogue of full-length 16S rRNA gene sequences and taxonomy for studies of bacterial communities in wastewater treatment plants. *Nat Commun* **13**, 1908 (2022).
3. Wu, L. *et al.* Global diversity and biogeography of bacterial communities in wastewater treatment plants. *Nature Microbiology* **4**, 1183–1195 (2019).
4. Jiang, C. *et al.* Characterizing the growing microorganisms at species level in 46 anaerobic digesters at Danish wastewater treatment plants: A six-year survey on microbial community structure and key drivers. *Water Research* **193**, 116871 (2021).
